# Proteasomal degradation of human SERINC4: a potent host anti-HIV-1 factor that is antagonized by Nef

**DOI:** 10.1101/2020.12.07.414888

**Authors:** Xusheng Qiu, Ifeanyichukwu E. Eke, Silas F. Johnson, Chan Ding, Yong-Hui Zheng

## Abstract

The serine incorporator (SERINC) protein family has five paralogous members with 9-11 transmembrane domains. SERINC5 is a potent host restriction factor and antagonized by HIV-1 Nef and two other retroviral accessory proteins via the lysosomal degradation pathway. Here, we investigated human SERINC4 expression and antiviral mechanisms. Unlike its four paralogs, human SERINC4 is subjected to proteasome-mediated turnover, resulting in ~250-fold lower expression than SERINC5. However, when expression was normalized, human SERINC4 restricted HIV-1 replication as effectively as SERINC5, and SERINC4 was also antagonized by Nef via the lysosomal pathway. Although SERINC4 proteins are conserved within primates or rodents, their N-terminal regions are highly variable across species. Interestingly, unlike human SERINC4, murine SERINC4 was stably expressed but had a very poor antiviral activity. We created stable SERINC4 chimeras by replacing the N-terminal region and found that the 1-34 and 35-92 amino acids determine SERINC4 antiviral activity or protein expression, respectively. Using these chimeras, we demonstrate that SERINC4 is incorporated into HIV-1 virions and restricts Tier 1 HIV-1 more effectively than Tier 3 HIV-1. Importantly, SERINC4 increases HIV-1 sensitivity to broadly neutralizing antibodies. Thus, human SERINC4 strongly restricts HIV-1 replication when it is overexpressed, which reflects a potential antiviral activity of this gene product under physiological conditions.

**Highlights:**

- Identification of another potent anti-HIV-1 host factor SERINC4 from the SERINC family
- Identification of two N-terminal domains that regulate SERINC4 expression and antiviral activity
- Understanding the natural degradation of human SERINC4 by the proteasomal pathway
- Understanding the important role of the lysosomal pathway in Nef antagonism of host restriction

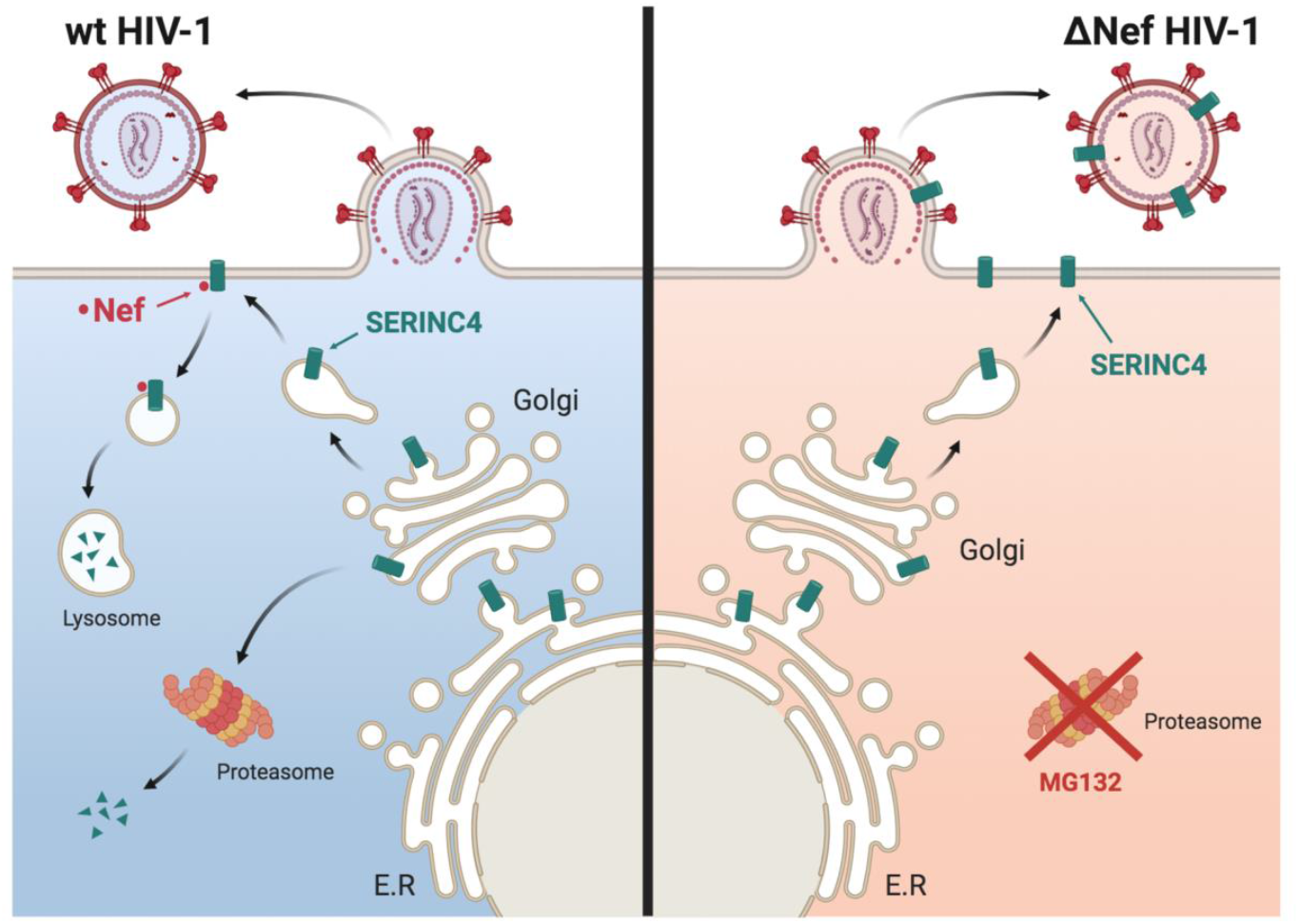

## Introduction

<u>Ser</u>ine <u>inc</u>orporator (SERINC) protein family was initially identified as serine transporters that was thought to play a role in phosphatidylserine and sphingolipid biosynthesis (Inuzuka et al., 2005). This SERINC (Ser) family has five members (Ser1 to Ser5) that are type III integral membrane proteins with 9-11 transmembrane (TM) domains and share 31-58% sequence homology (Inuzuka et al., 2005). Recently, Ser5 and Ser3 were identified as novel host restriction factors that are incorporated into HIV-1 virions and inhibit viral replication at virus entry (Rosa et al., 2015; Usami et al., 2015). Compared to Ser5, the Ser3 antiviral activity is very weak. The Ser5 antiviral activity is antagonized by HIV-1 Nef (Rosa et al., 2015; Usami et al., 2015), murine leukemia virus (MLV) glycosylated Gag (glycoGag) (Rosa et al., 2015; Usami et al., 2015), and equine infectious anemia virus (EIAV) S2 proteins (Ahi et al., 2016; Chande et al., 2016). We reported that Nef, glycoGag, and S2 proteins downregulate Ser5 from plasma membrane and target Ser5 to endosomes and lysosomes for degradation (Ahmad et al., 2019; Li et al., 2019; Shi et al., 2018). Thus, Ser5 is an important restriction factor for a wide range of retroviruses.

Ser5 inhibits virus entry at the stage of fusion pore formation after being incorporated into virions (Sood et al., 2017). Ser5 renders HIV-1 Env proteins more sensitive to broadly neutralizing antibodies (bNAbs) (Beitari et al., 2017; Lai et al., 2011; Sood et al., 2017; Usami and Gottlinger, 2013), suggesting that Ser5 modifies Env conformation by directly targeting those Env trimers. Indeed, the Ser5 antiviral activity is dependent on Env glycoproteins in a strain-specific manner. Tier 1 strains that are mostly laboratory-adapted viruses are sensitive, whereas the majority of Tier 2/3 viruses that are primary isolates are resistant to Ser5 restriction (Beitari et al., 2017; Sood et al., 2017; Usami et al., 2015). In fact, native Tier 1 Env trimers predominantly adopt a CD4-bound, open conformation, while Tier 2/3 Env trimers retain a pre-fusion, closed conformation (Munro et al., 2014; Munro and Mothes, 2015). We reported that Ser5 interacts with Env trimers in an open state more strongly and dissociate these open trimers, which may explain why Ser5 inhibits HIV-1 replication in an Env-dependent manner (Zhang et al., 2019).

Here, we investigated human Ser4 protein expression and anti-HIV-1 mechanisms by comparing this protein with its orthologs and paralogs. We found that human Ser4 is poorly expressed but has a strong antiviral activity. On the contrary, murine Ser4 is stably expressed but has a very poor antiviral activity. Via creating human and murine chimeric Ser4 proteins, we identified two separated N-terminal regions that differentially regulate Ser4 protein expression and its antiviral activity.

## Results

### Human Ser4 is poorly expressed but has a strong anti-HIV-1 activity

To compare levels of human Ser1, Ser2, Ser3, Ser4, and Ser5 expression, these proteins were tagged with a C-terminal FLAG epitope and expressed from pCMV6 mammalian expression vector. One microgram vectors were used to transfect 293T cells and their expression was detected by Western blotting (WB) using anti-FLAG. The expression of Ser1, Ser2, Ser3, and Ser5 was detected, but Ser4 expression was not (**Fig.1A**). To detect Ser4, undiluted Ser4 sample was analyzed again with serially diluted Ser5 samples. After a longer exposure, Ser4 expression was detected, but its signal intensity was only comparable to Ser5 that was diluted by ~256-fold (**Fig.1B**). Thus, human Ser4 is expressed at least 250-fold less than human Ser5 at steady-state levels.

**Figure 1.**
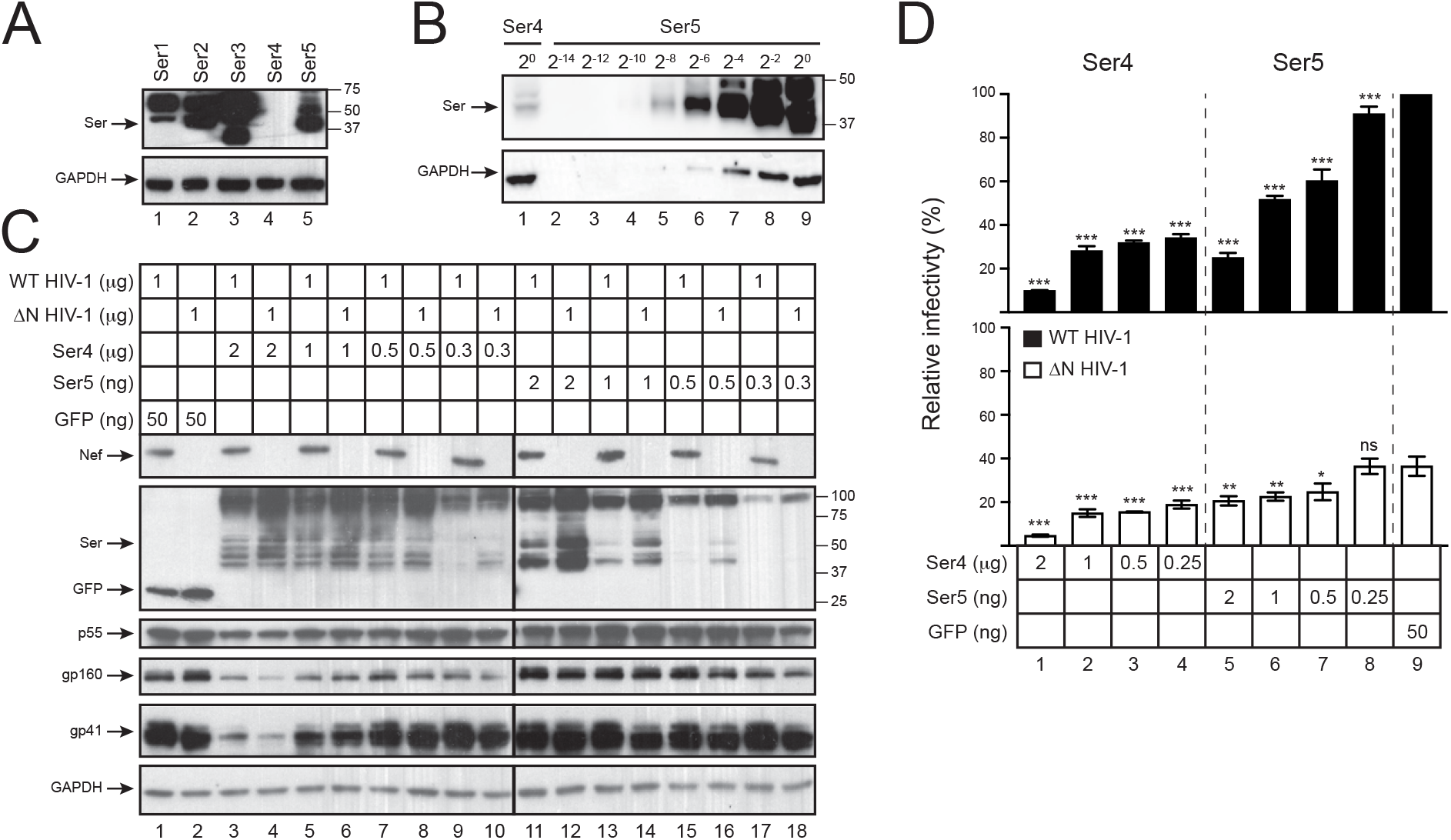
Analysis of human Ser4 expression and anti-HIV-1 activity. **(A)** 293T cells were transfected with 1 μg pCMV6 vectors expressing human Ser1, Ser2, Ser3, Ser4, or Ser5 that have a C-terminal FLAG tag. Protein expression was compared by WB using anti-FLAG, and GAPDH was used as loading controls. **(B)** The levels of Ser4 expression were compared to Ser5 by WB after serial dilutions. **(C)** Wild-type (WT) and *nef-deficient* (ΔN) NL4-3 viruses were produced from 293T cells after transfection with a pH22 or pH22ΔN HIV-1 proviral vector, in the presence of indicated amounts of pCMV6-Ser4, pCMV6-Ser5, or pcDNA-GFP that all express a C-terminal FLAG tag. Cellular Ser4, Ser5, and GFP expression were detected by anti-FLAG and cellular HIV-1 protein expression was detected by indicated antibodies via WB. **(D)** Viruses were collected from culture supernatants in (C). After normalization of viral titers by p24^Gag^ ELISA, viral infectivity was determined after infection of TZM-bI cells. Infectivity is shown as relative values, with the infectivity of WT viruses produced in the presence of GFP set as 100%. Error bars represent standard deviation (SDs) from three independent experiments. *p<0.05, **p<0.01, ***p<0.001, ns, not significant (p>0.05).

To detect Ser4 antiviral activity, wild-type (WT) and Nef-defective (ΔN) HIV-1 virions were produced from 293T cells in the presence of increasing amounts of Ser4 and Ser5 expression vectors using green fluorescent protein (GFP) as a negative control. To ensure that Ser4 and Ser5 were expressed at similar levels, 1,000-fold less Ser5 vectors were used for transfection. No difference in Ser4 protein expression was found when 2 μg, 1 μg, and 0.5 μg Ser4 vectors were used, indicating that the Ser4 expression was saturated under these conditions (**Fig.1C**, lanes 3 to 8). The Ser4 and Ser5 expression levels became comparable when 0.5 μg Ser4 and 2 ng Ser5 vectors were used for transfection (**Fig.1C**, lanes 7, 8, 11, 12), confirming that Ser4 is expressed ~250-fold less than Ser5. Although Ser4 reduced HIV-1 Gag and Env protein expression when 2 μg Ser4 vectors were used for transfection, neither Ser4 nor Ser5 affected the Gag and Env expression under the other transfection conditions. Thus, the reduction by Ser4 at 2 μg should be caused by a transfection artefact. In addition, levels of both Ser4 and Ser5 expression were reduced by Nef, indicating that Nef counteracts Ser4.

Next, virions were collected from these 293T cells and viral infectivity was determined via infection of the HIV-1 luciferase reporter TZM-bI cells. Consistently, both Ser4 and Ser5 reduced the ΔN virus infectivity much more strongly than WT virus infectivity in a dose-dependent manner, confirming that Ser4 is sensitive to Nef (**Fig.1D**). Importantly, both Ser4 and Ser5 reduced the ΔN HIV-1 infectivity to the same levels when they were expressed at a similar level in viral producer cells (Fig.1D, lanes 2 to 5). These results demonstrate that although Ser4 is poorly expressed, it has a similar level of anti-HIV-1 activity as Ser5 if the Ser4 expression can be upregulated.

### Human Ser4 proteins are targeted by proteasomes

To understand the poor Ser4 expression, we tested whether this protein is targeted by protein degradation pathways. When cells were treated with proteasome inhibitors (MG132, lactacystin) and lysosome inhibitors (NH4Cl, bafilomycin A1), the Ser4 expression was selectively increased by MG132 and lactacystin. In contrast, the Ser5 expression was slightly increased by both types of inhibitors except for MG132 (**Fig.2A**). Thus, unlike Ser5, human Ser4 is aggressively targeted to proteasomes for destruction.

**Figure 2.**
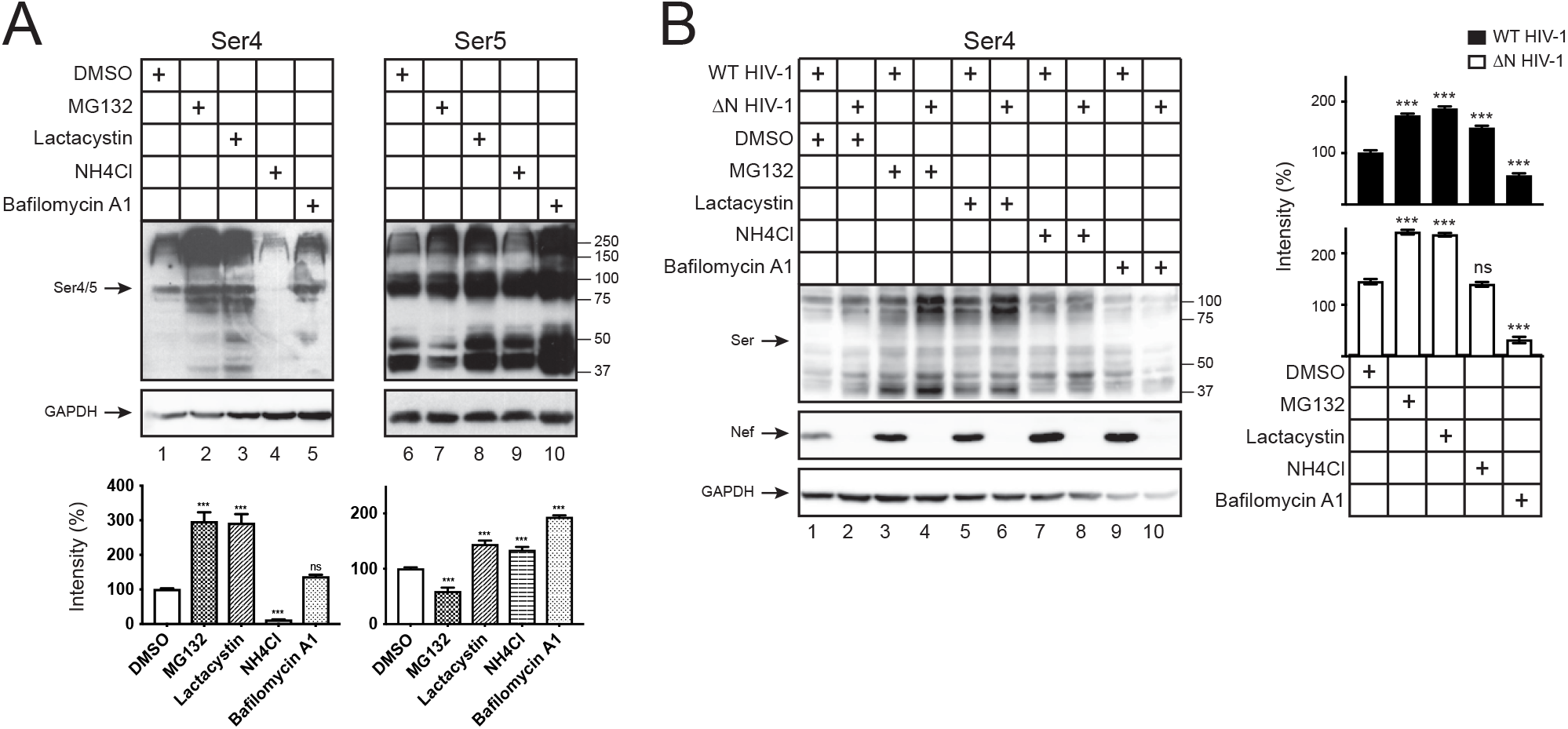
Human Ser4 is degraded in proteasomes. **(A)** 293T cells were transfected with 2 μg pCMV6-Ser4 or 0.5 μg pCMV6-Ser5. After treatments with DMSO (control), MG132 (10 μM), lactacystin (10 μM), NH4Cl (20 μM), or bafilomycin A1 (100 nM) for 12 h, Ser4 and Ser5 expression were determined by WB using anti-FLAG. The Ser4 or Ser5 expression levels were quantified using ImageJ and are presented as relative values. The levels of Ser4 and Ser5 samples treated with DMSO were set as 100%. **(B)** 293T cells were transfected with 1 μg pH22 or pH22ΔN that produces WT or ΔN NL4-3 viruses in the presence of 1 μg pCMV6-Ser4. After similar treatments as in (A), Ser4 and Ser5 expression were determined and quantified similarly as in (A). The levels of Ser4 in the presence of ΔN HIV-1 treated with DMSO were set as 100%. Error bars represent SDs from three independent experiments. *p<0.05, **p<0.01, ***p<0.001, ns, not significant (p>0.05).

We reported that HIV-1 Nef antagonizes Ser5 via the endosomes and lysosomes for degradation (Shi et al., 2018). To understand how Nef antagonizes Ser4, human Ser4 was expressed with WT and ΔN HIV-1 in 293T cells and treated with the same four inhibitors. Nef decreased the Ser4 expression by ~40% in the absence of any treatments (Fig.2B, lanes 1, 2), confirming Nef downregulation of Ser4. Again, MG132 and lactacystin increased the Ser4 expression (Fig.2B, lanes 3 to 6), whereas NH4Cl and bafilomycin did not (**Fig.2B**, lanes 7 to 10). Although Nef could still decrease the Ser4 expression in the presence of MG132 and lactacystin by ~70% (**Fig.2B**, lanes 3, 5), it became unable to do so when cells were treated with NH4Cl and bafilomycin A1. (Fig.2B, lanes 7, 9). These results demonstrate that Nef also employs the lysosomal pathway to antagonize Ser4.

### Murine Ser4 is stably expressed but has a very poor anti-HIV-1 activity

To know how Ser4 proteins from other species are expressed and whether they have any anti-HIV-1 activity, human and murine (m) Ser4 and Ser5 proteins were expressed with HIV-1 in 293T cells for viral production. Unlike human Ser4, the mSer4 expression was detected in these viral producer cells by WB, although its levels were still lower than human and murine Ser5 (**Fig.3A**). Levels of mSer4 expression were ~32-fold higher than human Ser4 (**Fig.3B**, lanes 8, 9). Thus, unlike human Ser4, mSer4 proteins are stably expressed. In addition, like mSer5, the mSer4 expression was also decreased by Nef (**Fig.3A**), indicating Ser4 from rodents is also sensitive to Nef.

**Figure 3.**
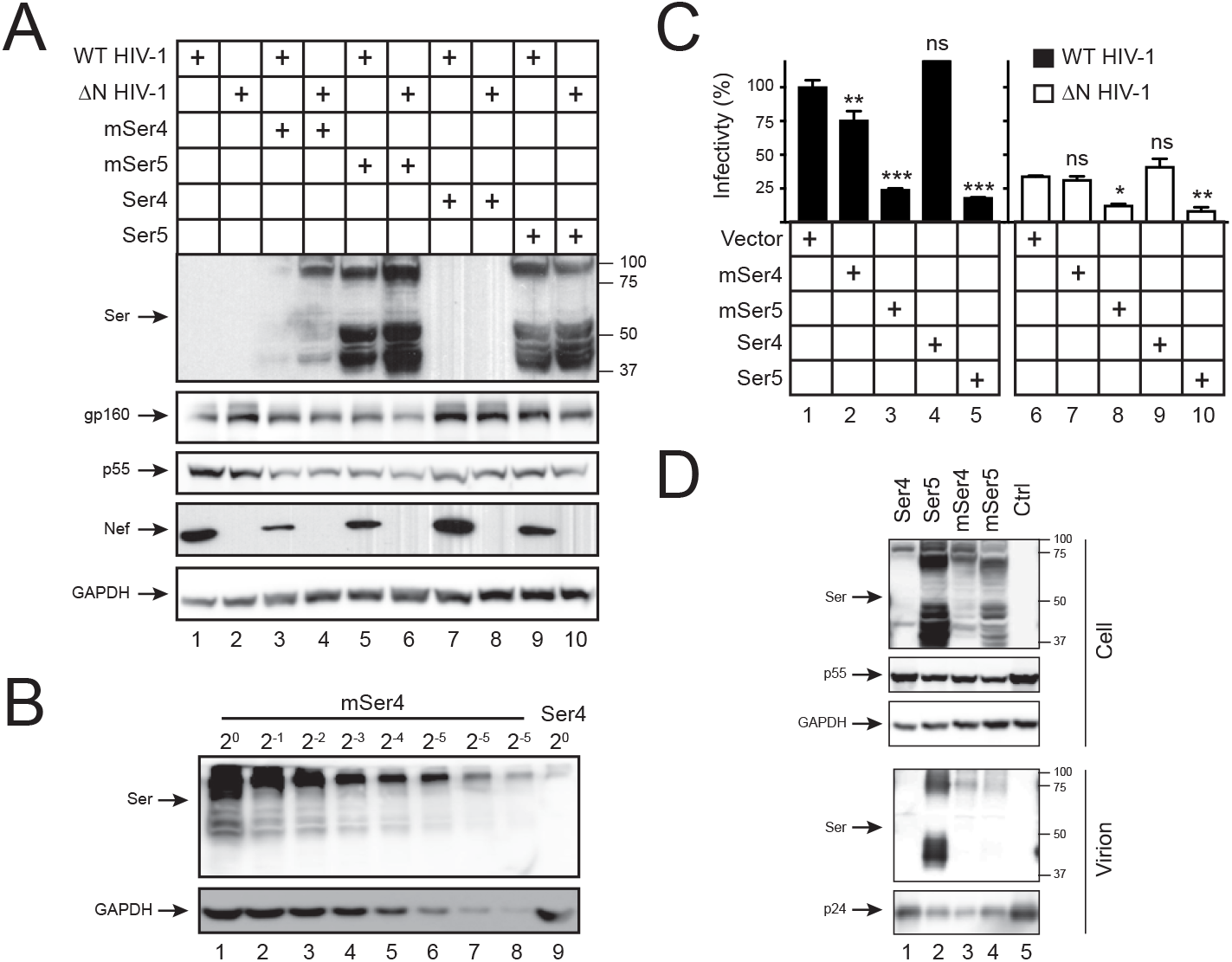
Analysis of mSer4 expression and anti-HIV-1 activity. **(A)** WT and ΔN HIV-1 NL4-3 viruses were produced from 293T cells in the presence of 100 ng pCMV6 vectors expressing human and murine Ser4 or Ser5. Cellular Ser4 and Ser5 expression were determined by anti-FLAG and cellular HIV-1 protein expression was detected by indicated antibodies via WB. **(B)** 293T cells were transfected with 1 μg pCMV6 vectors expressing mSer4 and human Ser4. mSer4-expressing cell lysate was serially diluted and compared to human Ser4 by WB using anti-FLAG. **(C)** Virions were collected from culture supernatants in (A), and viral infectivity was determined in TZM-bI cells. Infectivity is shown as relative values, with the infectivity of WT viruses produced in the presence of a control vector set as 100%. Error bars represent SDs from three independent experiments. *p<0.05, **p<0.01, ***p<0.001, ns, not significant (p>0.05).

We then analyzed the infectivity of virions produced from these 293T cells after infection of TZM-bI cells. Both human and murine Ser5 exhibited a very strong anti-HIV-1 activity (**Fig.3C**, lanes 3, 5, 8, 9). Due to the poor expression, human Ser4 did not show any antiviral activity (**Fig.3C**, lanes 4, 9). However, despite the high expression, mSer4 showed a similar low level of antiviral activity as human Ser4 (**Fig.3C**, lanes 2, 7). These results demonstrate that mSer4 barely restricts HIV-1 replication.

### The N-terminal 35-92 amino acids (aa) of human Ser4 determine its poor expression

To further explore the poor human Ser4 expression mechanism, we compared Ser4 protein sequences from primates and rodents. When human (h), gorilla (gor), macaque (mac), mouse (m) and rat Ser4 sequences were aligned, it is clear that Ser4 proteins are conserved within each species, but they differ significantly between primate and rodent species in both N-terminal and C-terminal regions (**Fig.S1**). Compared to primate Ser4 proteins, mSer4 has a large number of substitutions in the 1-92 aa region (**Fig.4A**). It was reported that human Ser5 has 10 TM domains (Pye et al., 2020). When the Ser4 topology was predicted by the transmembrane hidden Markov model (TMHMM), unlike mSer4, the likelihood for human Ser4 to have the 2^nd^ TM domain is significantly diminished (**Fig.4B**). Thus, we decided to focus on these 1-92 aa to study the Ser4 expression and antiviral mechanisms.

**Figure 4.**
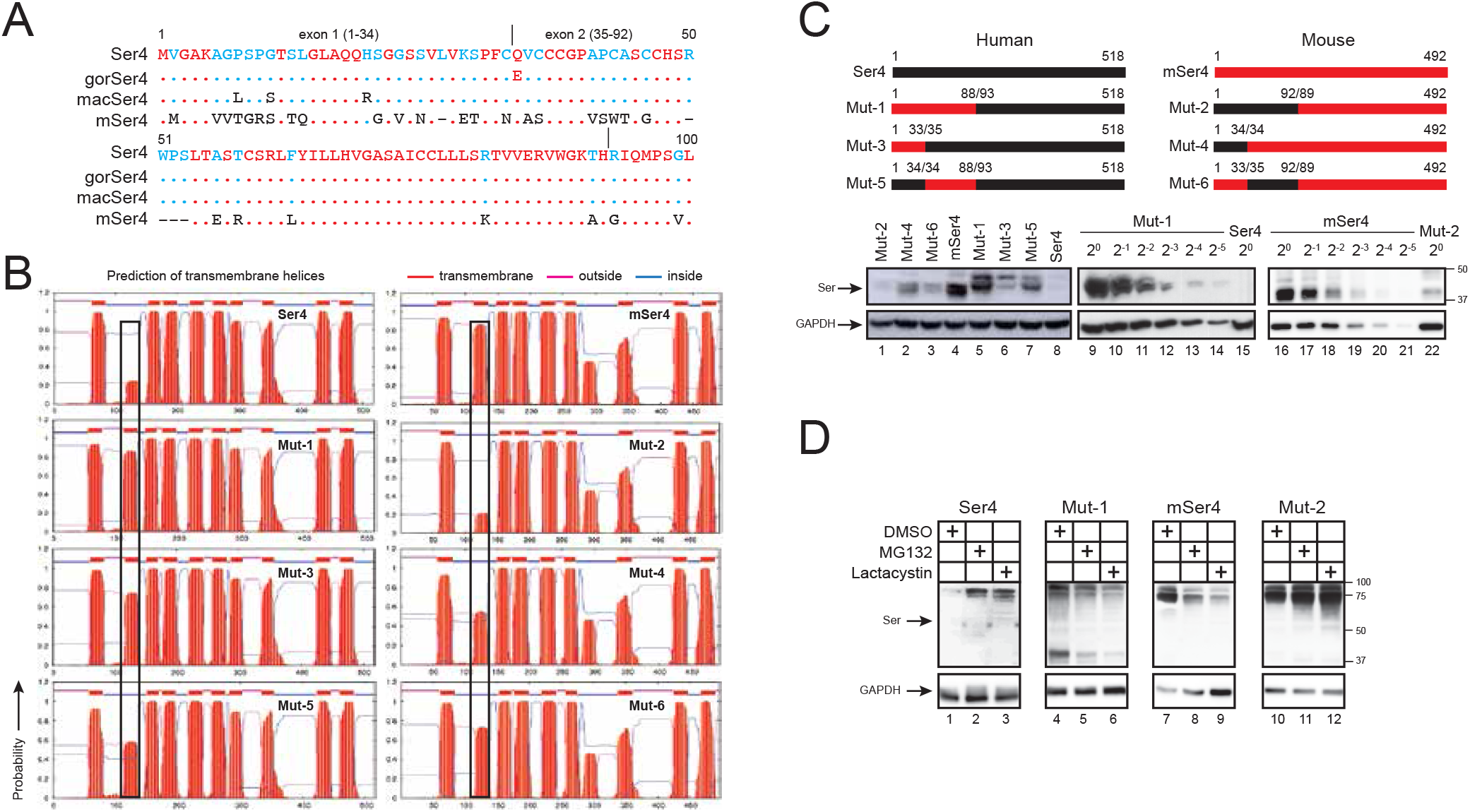
Analysis of human-mouse chimeric Ser4 expression. **(A)** The N-terminal amino acid sequences of human, gorilla (gor), macaque (mac), and murine (m) Ser4 proteins are aligned. Red, blue, black, and dash indicate conserved, partially conserved, non-conserved, or non-existing residues, respectively. Dots indicate identical residues. The exon 1 and exon 2 region are indicated. **(B)** Membrane topology of WT and chimeric Ser4 proteins were predicted from the public TMHMM server and are presented. The putative 2^nd^ TM domains are squared. **(C)** Chimeric Ser4 mutants (Mut-1, 2, 3, 4, 5, 6) were created by swapping the indicated human Ser4 (black) and murine Ser4 (red) region and expressed from the pCMV6 vector that expresses a C-terminal FLAG tag. 293T cells were transfected with 1 μg WT or these mutant expression vectors, and their expression was compared by WB using anti-FLAG. In addition, Mut-1 and mSer4-expressing cell lysate were serially diluted and compared to human Ser4 or Mut-2 by WB using anti-FLAG. **(D)** 293T cells were transfected with 1 μg pCMV6 vectors expressing human Ser4, mSer4, Mut-1, and Mut-2, and treated with DMSO (control), MG132 (10 μM), or lactacystin (10 μM) for 12 h. Their expression was determined by WB using anti-FLAG.

We created six Ser4 mutants by swapping the N-terminal regions between human and murine Ser4 (**Fig.4C**). Mut-1, Mut-3, and Mut-5 are human Ser4 proteins that express mSer4 1-88, 1-33, or 34-88 aa; Mut-2, Mut-4, and Mut-6 are mSer4 proteins that express human Ser4 1-92, 1-34, or 35-92 aa. Initially, we compared their transmembrane topologies. The likelihood for the 2^nd^ TM domain is increased in Mut-1, but decreased in Mut-2, suggesting that the 1-92 aa could be a determinant for the formation of the 2^nd^ TM domain (**Fig.4B**). In addition, because Mut-3 and Mut-5 display increased, whereas Mut-4 and Mut-6 display decreased likelihood for the 2^nd^ TM domain, both the 1-34 and 35-92 region should contribute to the formation of the 2^nd^ TM domain.

Next, we compared their protein expression by WB. Mut-1 was expressed at least 64-fold higher than human Ser4 (**Fig.4C**, lanes 14, 15), whereas Mut-2 was ~16-fold lower than mSer4 (**Fig.4C**, lanes 19, 22). In addition, Mut-4 and Mut-5 were expressed at higher levels than Mut-6 or Mut-3 (**Fig.4C**, lanes 2, 3, 6, 7). These results demonstrate the 1-92 aa of human Ser4 are critical for its poor protein expression, and in this region, 35-92 aa are more critical than 1-34 aa.

To understand whether the poorly expressed Mut-2 is degraded similarly as human Ser4, human Ser4, mSer4, Mut-1, and Mut-2 were expressed in 293 cells and treated with MG132 and lactacystin. Although both inhibitors did not increase Mut-1 and mSer4 expressions (**Fig.4D**, lanes 5, 6, 8, 9), they increased Mut-2 and human Ser4 expression (**Fig.4D**, lanes 2, 3, 11, 12). Thus, Mut-2 is also targeted to proteasomes for degradation, resulting in poor protein expression.

### The N-terminal 1-34 aa of human Ser4 determine its anti-HIV-1 activity

To understand the poor mSer4 antiviral activity, human Ser4, mSer4, Mut-1, Mut-2, Mut-3, Mut-4, Mut-5, and Mut-6 were expressed with HIV-1 in 293T cells. It was confirmed again that Mut-1 was expressed at higher levels than Mut-2, so was Mut-5 than Mut-3, and Mut-4 than Mut-2 in these viral producer cells (**Fig.5A**). Next, virions were collected and viral infectivity was analysed in TZM-bI cells. Among the four human Ser4 proteins, only Mut-5 consistently showed some levels of antiviral activity as WT human Ser4, and Mut-3 did not show any antiviral activity (**Fig.5B**, lanes 6, 7, 8, 9). Among the four murine Ser4 proteins, Mut-2 and in particular, Mut-4, showed a very strong antiviral activity; WT mSer4 showed a marginal antiviral activity; and Mut-6 did not show any antiviral activity (**Fig.5B**, lanes 2, 3, 4, 5). Because antiviral activity was detected from Mut-4 and Mut-5 that express human Ser4 1-34 aa, but not from Mut-3 and Mut-6 that lack these residues, we conclude that 1-34 aa are critical for the human Ser4 antiviral activity.

**Figure 5.**
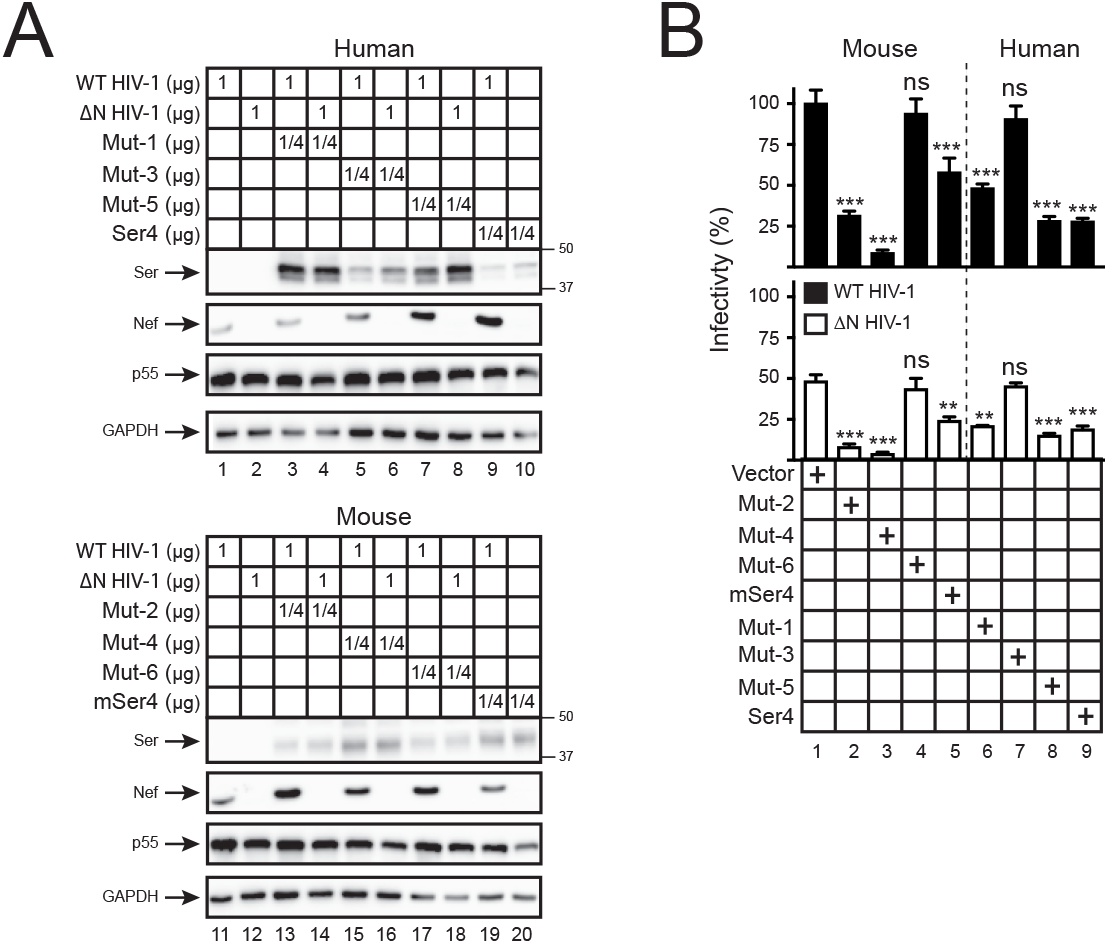
Analysis of human-mouse chimeric Ser4 anti-HIV-1 activity. **(A)** WT and ΔN HIV-1 NL4-3 viruses were produced from 293T cells in the presence of indicating amounts of pCMV6 vectors expressing Ser4, mSer4, and Mut-1 to 6. The expression of cellular Ser4 and its mutants was determined by anti-FLAG and that of cellular GAPDH and HIV-1 Nef and Gag was determined by specific antibodies via WB. **(B)** Viruses were collected from culture supernatants in (A), and viral infectivity was determined in TZM-bI cells. Infectivity is shown as relative values, with the infectivity of WT viruses produced in the presence of a control vector set as 100%. Error bars represent SDs from three independent experiments. *p<0.05, **p<0.01, ***p<0.001, ns, not significant (p>0.05).

### Rescue of human Ser4 expression by the human Ser2 and Ser5 N-terminal regions

Having found that the N-terminal region is responsible for the poor human Ser4 expression, we wondered whether its expression could be rescued by similar regions from the other human SERINC proteins. Two more mutants were created: Mut-7 that has the corresponding human Ser4 N-terminal amino acids replaced with the human Ser2 1-126 aa, and Mut-8 with the human Ser5 1-119 aa. Both Mut-7 and Mut-8 showed much higher likelihood for formation of the 2^nd^ TM domain than human Ser4 (**Fig.6A**). Consistently, unlike human Ser4, both Mut-7 and Mut-8 expression were detected by WB (**Fig.6B**, lanes 3, 5). Levels of Mut-7 expression were much higher than Mut-8, which could be due to the higher Ser2 expression than Ser5. These results demonstrate that the human Ser4 expression can also be rescued by human Ser2 and Ser5 N-terminal regions.

**Figure 6.**
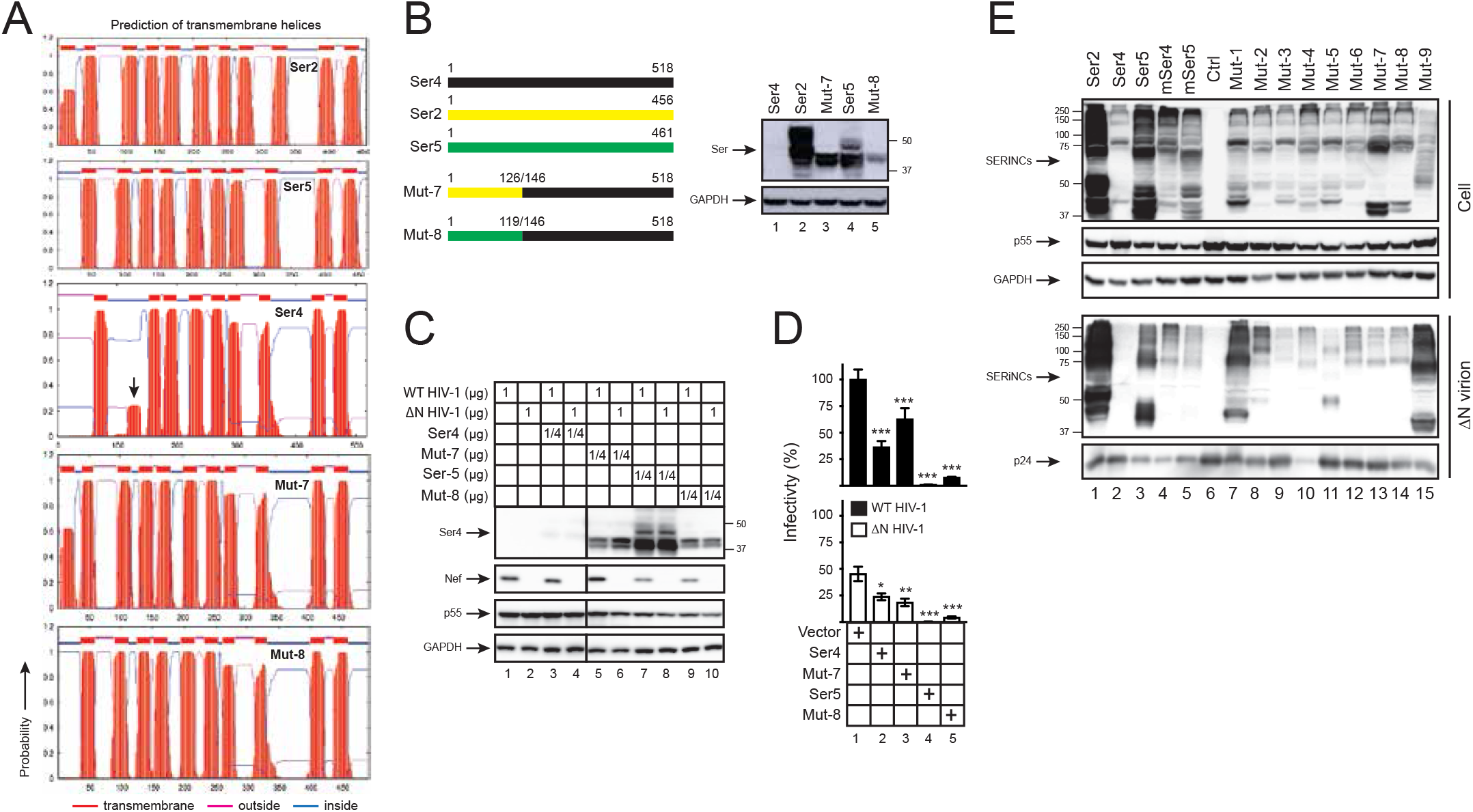
Analysis of Ser2-Ser4 and Ser5-Ser4 chimeric protein expression and anti-HIV-1 activity. **(A)** Ser2, Ser4, Ser5, and their chimeric protein topology were predicted by TMHMM and are presented. The putative human Ser4 2^nd^ TD domain is indicated by an arrow-head. **(B)** Mut-7 and Mut-8 were created by swapping the indicated human Ser4 (black) region with that from human Ser2 (yellow) or Ser5 (green), respectively, and expressed from the pCMV6 vector that expresses a C-terminal FLAG tag. 293T cells were transfected with 1 μg vectors expressing WT and these two mutants, and their expression was compared by WB using anti-FLAG. **(C)** 293T cells were transfected with pH22 or pH22ΔN that produces WT or ΔN NL4-3 viruses in the presence of indicating amounts of pCMV6 vectors expressing indicated Ser proteins. Cellular Ser protein expression was determined by anti-FLAG and cellular HIV-1 protein expression was determined by indicated antibodies via WB. **(D)** Viruses were collected from culture supernatants in (C), and viral infectivity was determined in TZM-bI cells. Infectivity is shown as relative values, with the infectivity of WT viruses produced in the presence of a control vector set as 100%. Error bars represent SDs from three independent experiments. *p<0.05, **p<0.01, ***p<0.001, ns, not significant (p>0.05). **(E)** 293T cells were cultured in 10-cm plates and transfected with 4.5 μg pH22ΔN and 4.5 μg pCMV6 vectors expressing indicated proteins. Virions were purified from culture supernatants via ultra-centrifugation and their expression in cells and virions was determined by WB.

Next, we measured the Mut-7 and Mut-8 antiviral activity. Again, the expression of human Ser5, Mut-7, and Mut-8, but not WT Ser4, was detected in viral producer cells by WB (**Fig.6C**). In target cells, Ser5 showed the strongest antiviral activity. Despite the relatively low-level expression, Mut-8 also showed a strong antiviral activity; and both WT Ser4 and Mut-7 showed a very marginal antiviral activity (**Fig.6D**). Collectively, these results demonstrate that only the Ser5, but not the Ser2 N-terminal region retains the human Ser4 anti-HIV-1 activity, which is consistent with that Ser2 does not have any anti-HIV-l activity.

### The Ser4 N-terminal region does not determine Ser4 packaging into virions

To further understand the poor Ser4 antiviral activity, we determined how these different Ser4 proteins are packaged into HIV-1 virions. ΔN HIV-1 virions were produced from 293T cells in the presence of human Ser2, Ser4, and Ser5, murine Ser4 and Ser5, and those eight Ser4 chimeras. Virions were purified by ultracentrifugation and SERINC protein expression was compared side-by-side by WB. Human Ser4 was not detected in virions, which was due to its low expression levels in cells (**Fig.6E**, lane 2). Human Ser2 and Ser5 and murine Ser4 and Ser5 were all detected in virions, and their levels of incorporation were proportional to their levels in cells, indicating a similar incorporation efficiency. All eight Ser4 chimeras were also detected in virions, indicating that their packing is not affected by the N-terminal domain.

### Ser4 restricts HIV-1 replication in an Env-dependent manner

In contrast to the poor human Ser4 expression and the poor mSer4 antiviral activity, Mut-4 and Mut-8 are expressed relatively well and show a strong antiviral activity. Thus, we used Mut-4 and Mut-8 to continue to study the Ser4 antiviral mechanism.

First, we compared their antiviral activity against HIV-1 Tier 1 NL4-3 and Tier 3 AD8 viruses using human Ser5 as a control. As reported previously (Beitari et al., 2017), Ser5 reduced the infectivity of NL4-3 much more effectively than AD8 in a dose-dependent manner (**Fig.7A**). Notably, so did Mut-4 and Mut-8, suggesting that NL4-3 viruses are also sensitive, whereas AD8 viruses are also resistant to Mut-4 and Mut-8. Thus, Ser4 shares the same Env-dependent antiviral mechanism as Ser5.

**Figure 7.**
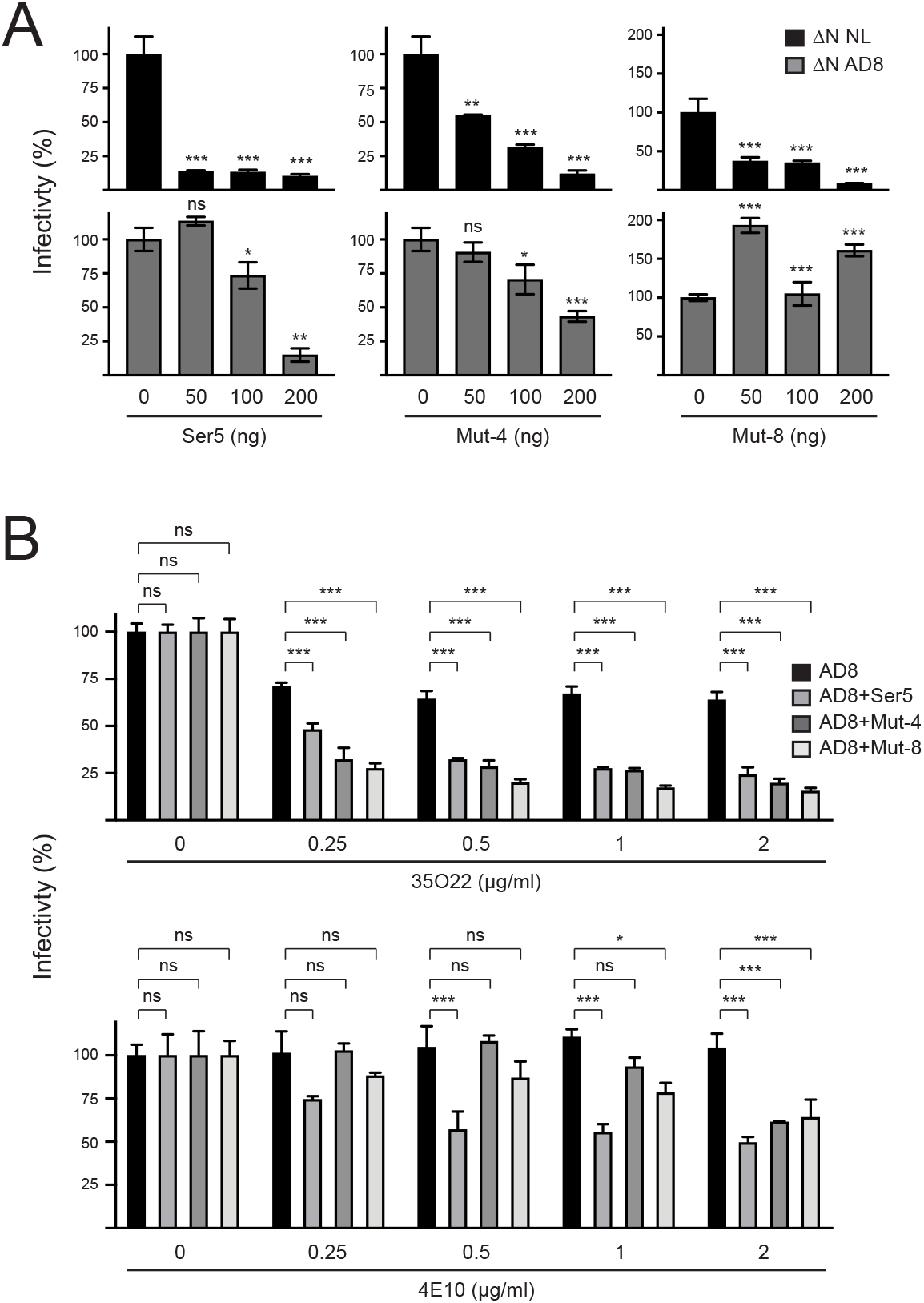
Analysis of Ser4 antiviral mechanism. **(A)** HIV-1 ΔN NL4-3 and ΔN AD8 viruses were produced from 293T cells after transfection with pH22ΔN or pH-AD8ΔN in the presence of indicated amounts of pCMV6 vectors expressing human Ser5, Mut-4, or Mut-8, and viral infectivity was determined in TZM-bI cells. The infectivity was presented as relative values, with that produced in the presence of a control vector set as 100%. **(B)** HIV-1 ΔN AD8 viruses were produced from 293T cells after transfection with pH-AD8ΔN in the presence of pCMV6-Ser5, pCMV6-Mut-4, or pCMV6-Mut-8. Viruses were treated with indicated amounts of HIV-1 bNAbs 35O22 or 4E10 at 37 °C for 1 h, and viral infectivity was determined after infection of TZM-bI cells. The infectivity was presented as relative values, with that of untreated viruses produced in the presence of a control vector set as 100%. Error bars in (A) and (B) represent SDs from three independent experiments. *p<0.05, **p<0.01, ***p<0.001, ns, not significant (p>0.05).

Second, we tested how these two chimeras affect HIV-1 sensitivity to bNAbs. AD8 viruses were produced in the presence of Ser5, Mut-4, or Mut-8. After incubation with two HIV-1 bNAbs including 35O22 and 4E10, viral infectivity was determined by infection of TZM-bI cells. Both bNAbs effectively reduced AD8 infectivity in a dose-dependent manner only in the presence of Ser5, Mut-4, and Mut-8. Thus, like Ser5, both Mut-4 and Mut-8 also increase HIV-1 sensitivity to bNAbs.

## Discussion

We found that human Ser4 is targeted to proteasomes for degradation, resulting in ~250-fold lower expression than human Ser5. In order to demonstrate the Ser4 antiviral activity, the Ser4 expression was normalized to the same level of Ser5 by overexpression. Under such condition, both Ser4 and Ser5 had a similar level of anti-HIV-1 activity. In addition, when the Ser4 expression was stabilized by replacing its N-terminal region, a strong Ser4 antiviral activity could also be detected. The Ser4 antiviral activity was also observed by Schulte *et al.* (Schulte et al., 2018). They overcome the poor human Ser4 expression by codon optimization, resulting in reduction of HIV-1 infectivity by ~50-fold. Thus, although human SERINC4 is unstable, its strong anti-HIV-1 activity can be detected by overexpression, indicating a potential antiviral activity of this gene product under physiological conditions.

The predicted Ser4 protein molecular weight is ~57 kDa. Nonetheless, Schulte *et al.* detected monomeric Ser4 proteins at ~45-kDa by SDS-PAGE. In addition, major Ser4 species were detected at high-molecular-weight (HMW) from 50-kDa to 100-kDa, which were preferentially packaged into HIV-1 virions (Schulte et al., 2018). We also detected these monomeric and HMW Ser4 species in cells, and the HMW species in HIV-1 virions. It is possible that these HMW Ser4 species were generated from protein aggregation, which frequently occurs during sample preparation of membrane proteins during WB. However, because these HMW species were selectively detected in virions, a more specific mechanism should be considered. It was reported that *N*-givcosviation contributes to the formation of HMW Ser5 species that are also preferentially packaged into virions (Sharma et al., 2018). Thus, it should be further investigated how Ser4 proteins are post-translationally modified and how the modification contributes to its function.

Among those ten Ser5 TM domains, the 10^th^ TM domain is required for Ser5 stable expression and antiviral activity (Zhang et al., 2017). In addition, the region between the 4^th^ and 8^th^ TM domain is required for Ser5 sub-cellular localization, virion incorporation, and antiviral activities (Schulte et al., 2018). It appears that regions important for Ser4 to restrict HIV-1 replication are distinct from those reported for Ser5. In the human Ser4 N-terminal region, we found that the 1-34 aa from exon 1 determines Ser4 antiviral activity, and the 35-92 aa from exon 2 determines Ser4 expression.

Our topology prediction indicates that the 2^nd^ TM domain of human Ser4 is unlikely to exist due to its N-terminal region. If this is indeed the case, the topology of human Ser4 should be different from the 2^nd^ TM domain onwards, resulting in a quite different protein compared to the other stably expressed proteins. Human Ser4 may have a very different glycosylation profile. It was reported that glycosylation-deficient Ser5 proteins are targeted to proteasomes for degradation, suggesting that glycosylation is important for Ser5 protein stability (Sharma et al., 2018). These results reiterate the importance of post-translational modification in human Ser4 expression. In addition, our prediction also suggests that Ser2 has an extra N-terminal TM domain that was absent in the previous report (Schulte et al., 2018). Like Ser2, Mut-7 also carries this extra TM domain and it is also well expressed. However, both Ser2 and Mut-2 have a very poor antiviral activity. Thus, it will be interesting to continue to investigate how the N-terminal TM domain contributes antiviral activity.

We found that both human and murine Ser4 expression were reduced by Nef, indicating that Nef counteracts Ser4. We reported that the Nef antagonism of Ser5 is easily saturable by Ser5 overexpression (Zhang et al., 2017). Consistently, the Nef antagonism was detected by WB when low levels of Ser5 were expressed, but was not detected when higher levels of Ser5 were expressed. Despite the very aggressive proteasomal targeting of human Ser4, Nef could still re-target it to the lysosomal pathway to antagonize its antiviral activity. Thus, Nef has evolved a very conservative mechanism to counteract SERINC proteins. We speculate that although the vast majority of human Ser4 proteins are eliminated by proteasomes during protein biosynthesis, residual human Ser4 proteins are survived and secreted to plasma membrane, which are incorporated into HIV-1 virions and restrict viral replication. Nonetheless, these cell surface Ser4 proteins are targeted by Nef, and delivered to endosomes and lysosomes for degradation.

## EXPERIMENTAL PROCEDURES

### Cells

Human 293T cells were obtained from ATCC. TZM-bI cells were obtained from NIH AIDS Reagent Program. These cells are cultured in DMEM with 10% bovine calf serum (HyClone).

### Plasmids

The HIV-1 proviral vectors pH22, pH22ΔN, pH-AD8, and pH-AD8ΔN were described before (Zhang et al., 2019). pCMV6 vectors expressing human Ser1, Ser2, Ser3, Ser4, and Ser5 and murine Ser4 that express a C-terminal FLAG tag were purchased from Origene, and the Ser1, Ser2, Ser3, and Ser5 expression and antiviral viral activity were reported in our previous publication (Zhang et al., 2017). To create Mut-1 to Mut-6, the exon 1 and exon 2 of human and murine Ser4 were amplified and ligated to the corresponding Ser4 by overlapping PCR. To create Mut-7 to Mut-8, regions covering the 1^st^, 2^nd^ and partial 3^rd^ TM domain were amplified from human Ser2 and Ser5 and ligated to human Ser4 via overlapping PCR. All these 8 mutants were cloned into pCMV6 vector after EcoRI/XhoI digestion, and they all express a C-terminal FLAG tag. Primers for cloning these 8 mutants are listed in supplemental Table 1. Their vector maps were created using SnapGene and are available upon request.

### HIV-1 production and infectivity analysis

To determine the antiviral activity of Ser4 and its mutants, viruses were produced from 293T cells after transfection with 1μg pH −22, pH22ΔN, pHAD8, pH-AD8ΔN together with various amounts of Ser4 expression vectors using polyethylenimine (PEI). Forty-eight hours later, the culture supernatants were collected, and the virus titer in which was quantified by p24^Gag^ ELISA as reported (Wehrly and Chesebro, 1997). To determine viral infectivity, equal amounts of viruses as normalized to the levels of p24^Gag^ were used to infect the HIV-1 luciferase reporter cell line TZM-bI in a 96-well plate at a density of 1 x 10^4^ per well. After 48 h, cells were lysed and intracellular luciferase activities were determined using Firefly Luciferase Assay Kit 2.0 (Biotium). These luciferase activities were used to calculate viral infectivity.

### HIV-1 neutralization assay

The HIV-1 broadly neutralizing antibodies 4E10 (catalog number 10091) (Stiegler et al., 2001) and 35O22 (catalog number 12586) (Huang et al., 2014) were obtained from the NIH AIDS Reagent Program. Prior to infection, viruses were incubated with serially diluted HIV-1 neutralizing antibodies at 37°C for 1 h. After that, viral infectivity was determined after infection of TZM-bI cells as described previously.

### Western blotting (WB)

Cells were lysed with 1% NP-40 in phosphate-buffered saline (PBS) containing protease inhibitor cocktail (Sigma). After removal of nucleus by low-speed centrifugation, samples were resolved by SDS PAGE gels. After transferring to Immun-Blot PVDF membrane (Bio-Rad), proteins were incubated with primary and secondary antibodies. The anti-Gag (Cat# 1513) and anti-gp41 (Cat# 526) were obtained from NIH AIDS Reagent Program. Mouse anti-glyceraldehyde-3-phosphate dehydrogenase (GAPDH) monoclonal antibody was purchased from Meridian Life Science. Horseradish peroxidase (HRP)-conjugated anti-human, rabbit, or mouse immunoglobulin G secondary antibodies were purchased from Pierce. HRP-conjugated anti-FLAG M2 antibody was purchased from Sigma. The enhanced chemiluminescence detection kit was purchased from Amersham Bioscience. Signals on Western blots were quantified by measuring protein band intensities using ImageJ (NIH, USA).

### Virion incorporation assay

To detect Ser protein incorporation into HIV-1 particles, 293T cells were cultured in 10-cm dishes and transfected with 6 μg HIV-1 proviral construct pH22ΔN and 3 μg Ser expression vectors. Supernatants were harvested 48 h after transfection and then centrifugated at 5,000 g for 10 min at 4°C to remove the cell debris. Virions were further purified by spinning the clarified supernatants through a 20% sucrose cushion at 296,000 g for 30 min at 4°C using the S100AT6 rotor (Sorvall). Pellets were dissolved in PBS and analyzed by WB.

## Statistics

Statistical tests were performed using GraphPad Prism 8. Variance was estimated by calculating the standard deviation (SD) and represented by error bars. Significance of differences between samples was assessed using two-way ANOVA with Bonferroni post-test. All experiments were performed independently at least three times, with a representative experiment being shown. *p<0.05, **p<0.01, ***p<0.001, ns, not significant (p>0.05).

## Acknowledgements

We thank NIH AIDS Reagent Program for providing various reagents.

## Supplemental Materials

**Supplemental Figure 1.**
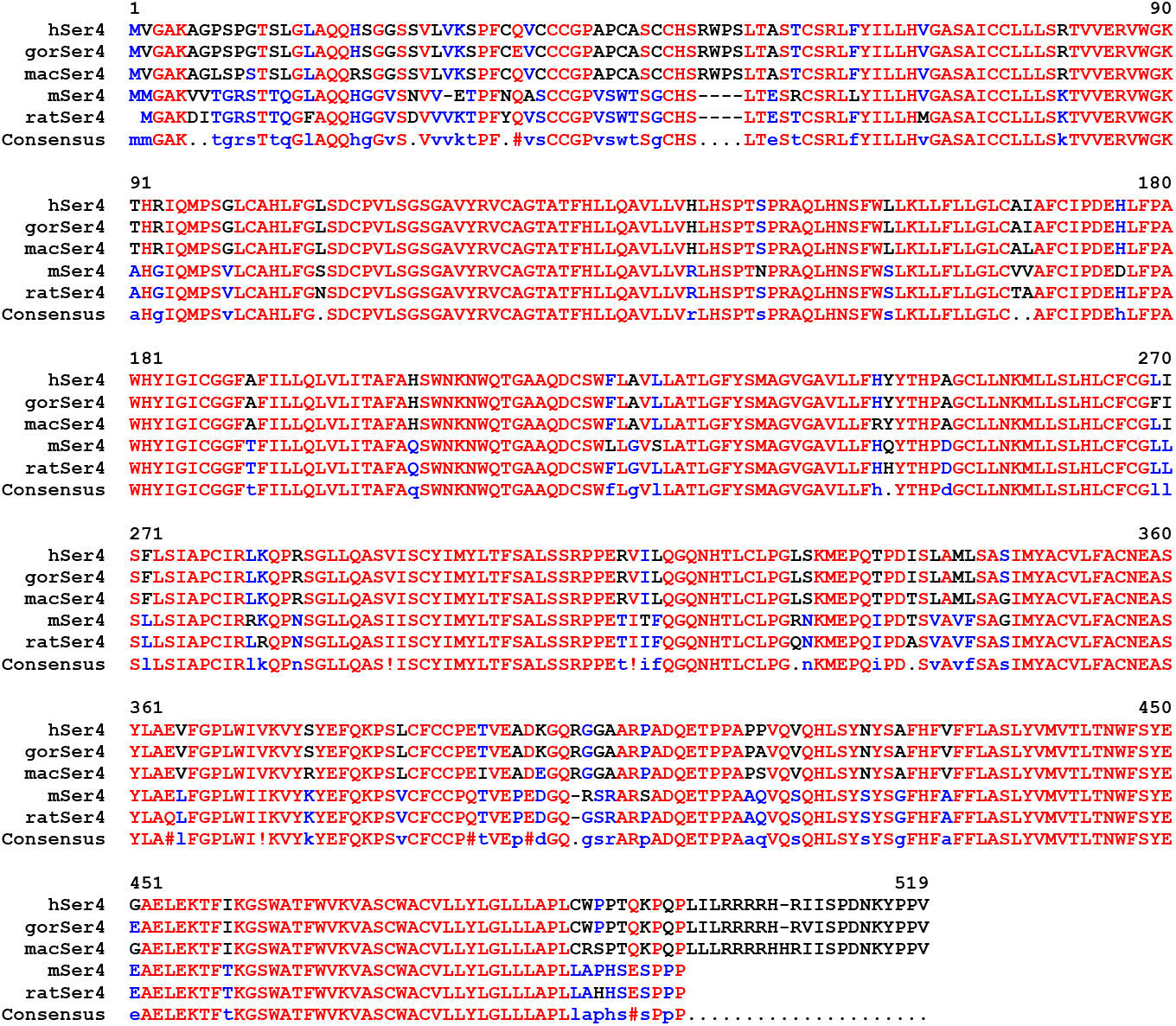
Ser4 amino acid sequences from human, gorilla (gor), macaque (mac), murine (m), and rat Ser4 proteins are aligned. Red, blue, black, and dash indicate conserved, partially conserved, non-conserved, or non-existing residues, respectively.

**Supplemental Table 1.**
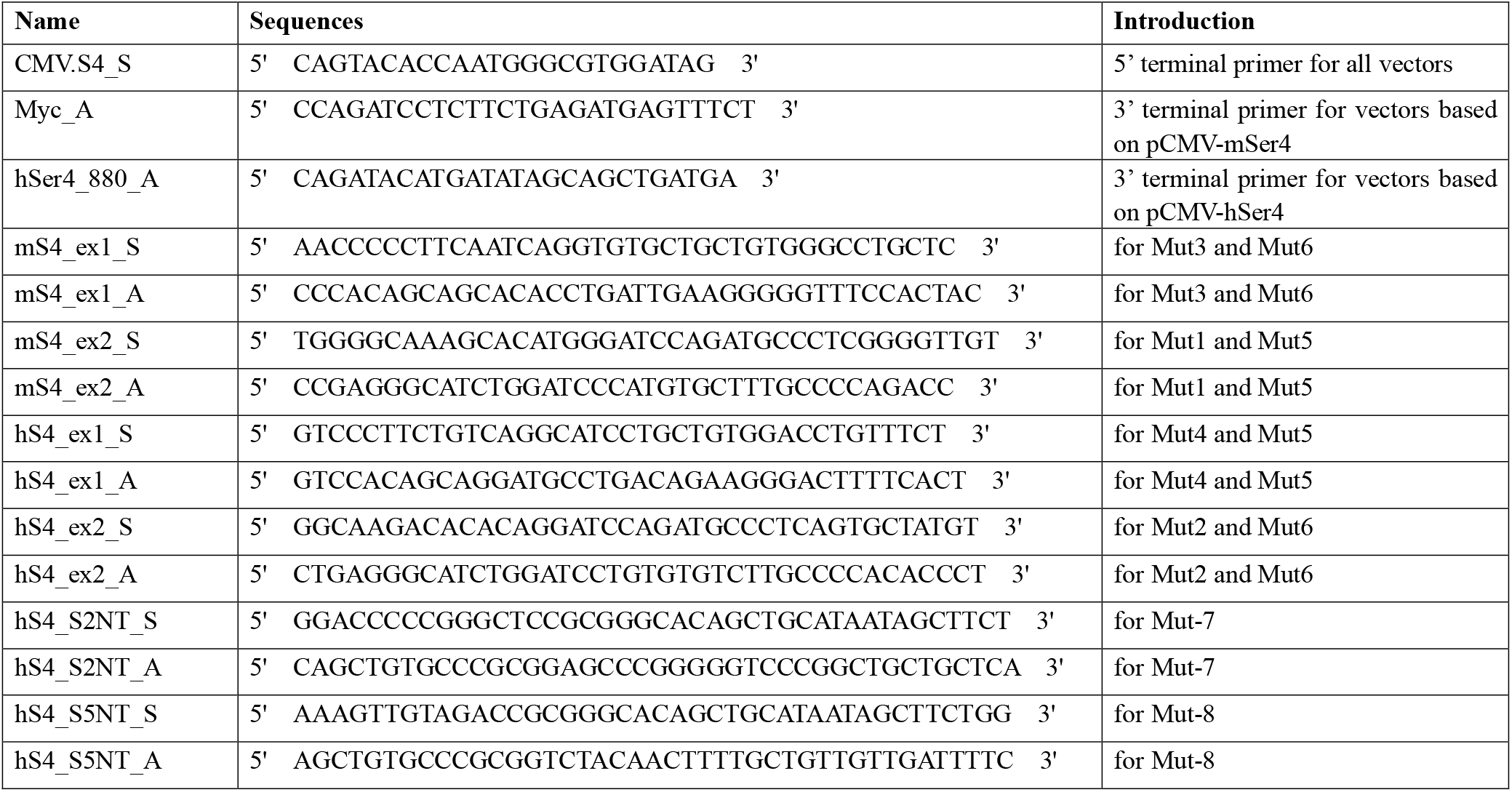
Primers used for creating Mut-1 to 8 by overlapping PCR are shown.

## References

Ahi, Y.S., Zhang, S., Thappeta, Y., Denman, A., Feizpour, A., Gummuluru, S., Reinhard, B., Muriaux, D., Fivash, M.J., and Rein, A. (2016). Functional Interplay Between Murine Leukemia Virus Glycogag, Serinc5, and Surface Glycoprotein Governs Virus Entry, with Opposite Effects on Gammaretroviral and Ebolavirus Glycoproteins. MBio 7.

Ahmad, I., Li, S., Li, R., Chai, Q., Zhang, L., Wang, B., Yu, C., and Zheng, Y.H. (2019). The retroviral accessory proteins S2, Nef, and glycoMA use similar mechanisms for antagonizing the host restriction factor SERINC5. J Biol Chem 294, 7013–7024.

Beitari, S., Ding, S., Pan, Q., Finzi, A., and Liang, C. (2017). Effect of HIV-1 Env on SERINC5 Antagonism. J Virol 91.

Chande, A., Cuccurullo, E.C., Rosa, A., Ziglio, S., Carpenter, S., and Pizzato, M. (2016). S2 from equine infectious anemia virus is an infectivity factor which counteracts the retroviral inhibitors SERINC5 and SERINC3. Proc Natl Acad Sci U S A 113, 13197–13202.

Huang, J., Kang, B.H., Pancera, M., Lee, J.H., Tong, T., Feng, Y., Imamichi, H., Georgiev, I.S., Chuang, G.Y., Druz, A., et al. (2014). Broad and potent HIV-1 neutralization by a human antibody that binds the gp41-gp120 interface. Nature 515, 138–142.

Inuzuka, M., Hayakawa, M., and Ingi, T. (2005). Serinc, an activity-regulated protein family, incorporates serine into membrane lipid synthesis. J Biol Chem 280, 35776–35783.

Lai, R.P., Yan, J., Heeney, J., McClure, M.O., Gottlinger, H., Luban, J., and Pizzato, M. (2011). Nef decreases HIV-1 sensitivity to neutralizing antibodies that target the membrane-proximal external region of TMgp41. PLoS Pathog 7, e1002442.

Li, S., Ahmad, I., Shi, J., Wang, B., Yu, C., Zhang, L., and Zheng, Y.H. (2019). Murine Leukemia Virus Glycosylated Gag Reduces Murine SERINC5 Protein Expression at Steady-State Levels via the Endosome/Lysosome Pathway to Counteract SERINC5 Antiretroviral Activity. J Virol 93.

Munro, J.B., Gorman, J., Ma, X., Zhou, Z., Arthos, J., Burton, D.R., Koff, W.C., Courter, J.R., Smith, A.B., 3rd, Kwong, P.D., et al. (2014). Conformational dynamics of single HIV-1 envelope trimers on the surface of native virions. Science 346, 759–763.

Munro, J.B., and Mothes, W. (2015). Structure and Dynamics of the Native HIV-1 Env Trimer. J Virol 89, 5752–5755.

Pye, V.E., Rosa, A., Bertelli, C., Struwe, W.B., Maslen, S.L., Corey, R., Liko, I., Hassall, M., Mattiuzzo, G., Ballandras-Colas, A., et al. (2020). A bipartite structural organization defines the SERINC family of HIV-1 restriction factors. Nat Struct Mol Biol 27, 78–83.

Rosa, A., Chande, A., Ziglio, S., De Sanctis, V., Bertorelli, R., Goh, S.L., McCauley, S.M., Nowosielska, A., Antonarakis, S.E., Luban, J., et al. (2015). HIV-1 Nef promotes infection by excluding SERINC5 from virion incorporation. Nature 526, 212–217.

Schulte, B., Selyutina, A., Opp, S., Herschhorn, A., Sodroski, J.G., Pizzato, M., and Diaz-Griffero, F. (2018). Localization to detergent-resistant membranes and HIV-1 core entry inhibition correlate with HIV-1 restriction by SERINC5. Virology 515, 52–65.

Sharma, S., Lewinski, M.K., and Guatelli, J. (2018). An N-Glycosylated Form of SERINC5 Is Specifically Incorporated into HIV-1 Virions. J Virol 92, e00753–00718.

Shi, J., Xiong, R., Zhou, T., Su, P., Zhang, X., Qiu, X., Li, H., Li, S., Yu, C., Wang, B., et al. (2018). HIV-1 Nef Antagonizes SERINC5 Restriction by Downregulation of SERINC5 via the Endosome/Lysosome System. J Virol 92.

Sood, C., Marin, M., Chande, A., Pizzato, M., and Melikyan, G.B. (2017). SERINC5 protein inhibits HIV-1 fusion pore formation by promoting functional inactivation of envelope glycoproteins. J Biol Chem 292, 6014–6026.

Stiegler, G., Kunert, R., Purtscher, M., Wolbank, S., Voglauer, R., Steindl, F., and Katinger, H. (2001). A potent cross-clade neutralizing human monoclonal antibody against a novel epitope on gp41 of human immunodeficiency virus type 1. AIDS Res Hum Retroviruses 17, 1757–1765.

Usami, Y., and Gottlinger, H. (2013). HIV-1 Nef responsiveness is determined by Env variable regions involved in trimer association and correlates with neutralization sensitivity. Cell Rep 5, 802–812.

Usami, Y., Wu, Y., and Gottlinger, H.G. (2015). SERINC3 and SERINC5 restrict HIV-1 infectivity and are counteracted by Nef. Nature 526, 218–223.

Wehrly, K., and Chesebro, B. (1997). p24 antigen capture assay for quantification of human immunodeficiency virus using readily available inexpensive reagents. Methods 12, 288–293.

Zhang, X., Shi, J., Qiu, X., Chai, Q., Frabutt, D.A., Schwartz, R.C., and Zheng, Y.H. (2019). CD4 Expression and Env Conformation Are Critical for HIV-1 Restriction by SERINC5. J Virol 93.

Zhang, X., Zhou, T., Yang, J., Lin, Y., Shi, J., Zhang, X., Frabutt, D.A., Zeng, X., Li, S., Venta, P.J., et al. (2017). Identification of SERINC5-001 as the Predominant Spliced Isoform for HIV-1 Restriction. J Virol 91.

